# Spectral signatures of cross-modal attentional control in the adolescent brain and their link with physical activity and aerobic fitness levels

**DOI:** 10.1101/2023.01.30.526274

**Authors:** Doris Hernández, Jan Kujala, Erkka Heinilä, Ilona Ruotsalainen, Hanna-Maija Lapinkero, Heidi Syväoja, Lauri Parkkonen, Tuija H. Tammelin, Tiina Parviainen

## Abstract

Top–down attentional control seems to increase and suppress the activity of sensory cortices for relevant stimuli and to suppress activity for irrelevant ones. Higher physical activity (PA) and aerobic fitness (AF) levels have been associated with improved attention, but most studies have focused on unimodal tasks (e.g., visual stimuli only). The impact of higher PA or AF levels on the ability of developing brains to focus on certain stimuli while ignoring distractions remains unknown. The aim of this study was to examine the neural processes in visual and auditory sensory cortices during a cross-modal attention–allocation task using magnetoencephalography in 13–16-year-old adolescents (*n* = 51). During continuous and simultaneous visual (15 Hz) and auditory (40 Hz) noise-tagging stimulation, participants attended to either visual or auditory targets appearing on their left or right sides. High and low PA groups were formed based on seven-day accelerometer measurements, and high and low AF groups were determined based on the 20-m shuttle-run test. Steady-state (evoked) responses to the visual stimulus were observed in all the adolescents in the primary visual cortex, but some did not show responses in the primary auditory cortices to the auditory stimulus. The adolescents with auditory-tag-driven signals in the left temporal cortex were older than those who did not show responses. Visual cortices showed enhanced visual-tag-related activity with attention, but there was no cross-modal effect, perhaps due to the developmental effect observed in the temporal areas. The visual-tag-related responses in the occipital cortex were enhanced in the higher-PA group, irrespective of task demands. In summary, sensory cortices are unequally involved in cross-modal attention in the adolescent brain. This involvement seems to be enhanced by attention. Higher PA seems to be associated with a specific visual engagement benefit in the adolescent brain.

**Highlights:**

- Visual and auditory cortices’ engagement differs in cross-modal processing in adolescence.
- Adolescents with responses in the left temporal cortex are older than those without responses.
- Physical activity, but not aerobic fitness, is associated with visual engagement benefits in the adolescent brain.

## 1. Introduction

We are confronted with huge amounts of sensory information in our daily lives. Attention is the cognitive capacity that allows us to focus on the relevant aspects of that information, while the irrelevant content is ignored. Our capacity for attention can be controlled or guided by goals or task requirements, namely top–down cognitive control or executive control (Diamond, 2013). Cognitive control is a key skill needed for any other cognitive competence (so-called central function). Top– down influence seems to enhance the activity in sensory cortices for items that are relevant and suppress activity for items irrelevant to task goals (Gazzaley & Nobre, 2012). Cognitive control develops rather late in children and adolescents (Davidson et al., 2006; Luna, 2009) and is therefore likely to be prone to different endogenous or external influences. Indeed, cognitive control of attention can be modulated by several factors such as emotions in a threatening situation (Dennis & Chen, 2007), other cognitive functions such as working memory resources (Myers et al., 2017), or the environment such as in the case of bilingual language switching in their native language or second language environment (Zhang et al., 2021). The different brain mechanisms and factors that might contribute to enhancing the capacity for attentional control remain to be fully understood.

Usually, unimodal paradigms (i.e., requiring the processing of only one sensory modality) have been used in the study of attention. Unimodal tasks are easy to use in lab settings, but they do not represent real-life situations well, which weakens their validity. A way forward to overcome this limitation would be to use cross-modal attention paradigms, where attention needs to be switched between two modalities (e.g., visual and auditory). Attention often occurs simultaneously through multiple sensory modalities, such as visual, auditory, and/or tactile. While each sensory cortex is designed to process a specific type of information, there seems to be considerable overlap in their representations between them (Karim et al., 2021; Rapp & Hendel, 2003). There is growing evidence suggesting that the processing of stimuli in one modality can be enhanced (Dematte et al., 2006; Zhao et al., 2021) or diminished (Geangu et al., 2021; Robinson et al., 2020) by the simultaneous presentation of stimuli in different modalities. Therefore, besides the well-established cortical areas that are suggested to provide “central attentional control” (i.e., prefrontal areas, cingulate gyrus, and posterior parietal cortex [Lepsien & Nobre, 2006]), it is possible that attentional demands in cross-modal tasks also modulate the sensory processing of information (Lage-Castellanos et al., 2022; van Atteveldt et al., 2014).

Interestingly, it seems that attention is one of the cognitive functions that is also influenced by changes in body physiology, namely by physical activity/exercise. The positive effects of physical activity/exercise on academic achievement (Alvarez-Bueno et al., 2017) have been speculated to stem from effects in cognitive functions, specifically attention/inhibition (Becker et al., 2014; Hillman et al., 2009). Physical exercise generates metabolic changes in the body that have been suggested to give rise to long-term modifications in the brain, both functionally and structurally (Cotman et al., 2007). Physical activity has also been linked to some effects on cognition, especially attention (Singh et al., 2019; Syväoja et al., 2014) and inhibitory control (Alvarez-Bueno et al., 2017; Wu et al., 2022). However, the “full pathway” of how the metabolic changes in the brain induced by higher involvement in regular physical activity can be linked to the observed changes in cognition has not been clearly shown. One confusing factor in the earlier literature is that physical activity (PA) and aerobic fitness (AF) used to be treated as comparable concepts. PA is a measure of the amount of movement done over time (Caspersen et al., 1985), while AF is considered a body condition people have or achieve because of physical activity, and they might well have different associations with brain structure and function.

To better understand the specific associations that either PA or AF levels have with brain structure and function, the same group of subjects needs to be used for both comparisons. Supporting the divergent role of PA vs. AF for the brain, earlier studies have indicated that AF is associated more strongly with brain structural measures than PA (Ruotsalainen et al., 2019, 2020), but PA might influence brain functional properties, which underlie cognition in adolescents (Hernández et al., 2021; Ruotsalainen et al., 2021).

In the domain of attention, we showed that a higher level of PA was associated with stronger modulation of brain oscillations underlying attentional control in adolescents, but AF did not show the same relationship (Hernández et al., 2021). Participants with higher levels of PA showed more reliance on spatial cues (either intentionally or implicitly) than adolescents with lower PA in a selective visual attention task. This association was mediated by the related interhemispheric asymmetry of alpha cortical oscillations. To move closer to real-life task settings, it is important to test whether similar PA-related brain functional modifications underlie cross-modal attentional capacities.

Some behavioral studies testing the associations of PA and AF with multisensory integration have used cross-modal task requirements. They showed that higher levels of acute and regular PA were associated with improvements in multisensory (visual–auditory) integration tasks in children (O’Brien et al., 2021) and older adults (Mahoney et al., 2015; Merriman et al., 2015; O’Brien et al., 2017). Interestingly, this is in line with the idea that at the time of maturational/aging-related modifications in the synaptic connections in the brain, it may be beneficial to engage in physical exercise to improve or maintain functional capacity.

To explore the possible role of PA and AF levels on cross-modal attention at the neural level in adolescents, we used a cross-modal attention paradigm with simultaneous recording of the neuromagnetic activity of the whole brain. The neural steady-state responses (SSRs) in the visual and auditory cortices during a cross-modal attention task provide a useful paradigm for separating the activation engaged in visual and auditory stimulation during attended or unattended conditions. The engagement of sensory cortices in task performance can be reliably followed by using a cross-modal attention paradigm, where each sensory cortex is probed by a continuous signal that contains a specific tag frequency. In the case of an audio–visual cross-modal attention paradigm, simultaneously presented continuous auditory and visual signals are amplitude modulated with a unique frequency. The modality-specific responses can later be extracted from the neural signal and contrasted between attended and unattended conditions. Due to its good spatial resolution, magnetoencephalography (MEG) allows for the extraction of modality-specific functional markers of attention allocation in the visual and auditory cortical areas.

First, we studied the cortical basis, specifically the role of sensory cortices, for task-relevant (attended) vs. task-irrelevant (unattended) information processing during cross-modal attention tasks. Second, we studied the associations between these attention effects at the visual and auditory sensory cortices and the behavioral measures of top–down attentional and inhibitory control. Finally, we explored the link between PA and/or AF levels with sensory cortex engagement, as well as behavioral performance measures, in this cross-modal (visual and auditory) attention task in the adolescent brain. We expected that the ongoing activation of visual and auditory cortices would modulate engagement in a task-specific manner and would reflect the level of attention required by the cross-modal attention task. Based on previous findings (Hernández et al., 2021; Ruotsalainen et al., 2020, 2021) suggesting a stronger role of PA than AF for functional brain measures, we also hypothesized that PA level would specifically show an association with the efficiency of neural engagement of sensory cortices to task-relevant signals (hence, difference between attended vs. unattended condition). To test our hypotheses, the attended and unattended cortical SSRs in each sensory cortex were contrasted between groups with high vs. low PA/AF in typically developed adolescents. An indicator of the efficiency of task-related attention (contrasting cortical entrainment of SSRs during attended vs. unattended conditions in each sensory cortex) was correlated with behavioral performance.

## 2. Methods

### 2.1. Participants

Participants were recruited from the sample of a larger study measuring the behavioral associations of AF/regular PA with cognition (Finnish Schools on the Move Program, Joensuu et al., 2018; Syväoja et al., 2019). All participants without neurological disorders, use of medication that influenced the central nervous system, or major medical conditions were invited to participate. Of these, 54 adolescents volunteered to participate in the study. Three participants were excluded from further analysis due to the low quality of the MEG data. All subjects (*n* = 51, 17 males and 34 females) were Finnish native speakers. Adolescents, as well as their legal guardians, signed informed consent forms at the beginning of the study. All subjects had normal or corrected to normal vision and hearing. The study was approved by the Central Finland Healthcare District Ethical Committee to be conducted in accordance with the Declaration of Helsinki.

Handedness was assessed using the Edinburgh Handedness Inventory, and all participants were classified as right-handed. Self-reports about their stage of puberty with the Tanner Scale (Marshall & Tanner 1969; 1970) were used to measure the pubertal development in our sample.

Participants were divided into two groups by their level of AF, estimated using the shuttle-run test (Léger et al., 1988), where the participants ran between two lines at an increasing speed (see Section 2.2 for details). The number of minutes that the participant lasted until exhaustion was taken as the measure of AF, and the third tertile (66%) was used as a cut-off point to divide the sample into high or moderate-to-low (modlow) AF groups once the data were normalized by gender and age.

Participants were also divided into two groups based on their PA levels, estimated by accelerometer data that were recorded during seven consecutive days (see the next paragraph for more details). The third tertile (66%) of moderate-to-vigorous PA (MVPA, see Section 2.2 for details) was used as a cutoff point to divide the sample into high or modlow PA groups. Table 1 summarizes the demographic information and statistical differences for the final sample, divided into two groups based on AF/PA measures.

**Table 1.** Demographic information (mean ± SD) and statistical differences for the groups based on physical activity and aerobic fitness.

|  | Physical activity |  | Aerobic fitness |  |
| --- | --- | --- | --- | --- |
|  | high | modlow | high | modlow |
| **Age (months)** | 171.93 ± 6.7 | 176.35 ± 12.3 | 177.39 ± 11.4 | 173.44 ± 9.7 |
| **Pubertal stage** | 3.09 ± 0.8 | 3.54 ± 0.9 | 3.25 ± 0.9 | 3.60 ± 0.8 |
| **Physical activity (MV, min/day)** | **77.73 ± 13.4** | **39.34 ± 10.4\*\*\*** | 58.32 ± 21.4 | 45.50 ± 19.9 |
| **Aerobic fitness (20-m shuttle run test, min)** | **7.02 ± 1.9** | **5.29 ± 1.9\*** | **7.70 ± 1.3** | **4.21 ± 1.4\*\*\*** |

\* $p < 0.05$
\*\*\* $p < 0.001$
SD: standard deviation; modlow: moderate-to-low; MV: moderate-to-vigorous

### 2.2. PA/AF measures

AF was measured using a maximal shuttle-run test (Leger et al., 1988), a measure widely used to estimate a person’s maximum oxygen uptake (VO_2_max). The test was performed as described by Nupponen et al. (1999) and specified for the present data collection by Joensuu et al. (2018). Participants were instructed to run between two lines separated by 20 m. An audio signal indicated that the speed should be accelerated. The speeds at the first and second levels were 8.0 and 9.0 km/h, respectively. After the second level, the speed increased by 0.5 km/h per level. The duration of each level was one minute. The participants’ level of AF was indicated by the time they spent until they failed to reach the end lines in two consecutive tones.

PA was measured using triaxial accelerometers (ActiGraph, Pensacola, FL, USA; models GT3X + and wGT3X+). Accelerometers were worn by the participants for seven consecutive days. Participants were instructed to wear the accelerometer on their right hips during waking hours, except for water-related activities. The data were collected at a sampling frequency of 60 Hz and were standardly filtered. A valid measurement day consisted of at least 10 h of data. Activity counts were collected in 15-s epochs. Periods of at least 30 min of consecutive zero counts were considered non-wear time. A customized Visual Basic macro for Excel software was used for data reduction. Moderate-to-vigorous physical activity (MVPA) was calculated as a weighted-mean value of MVPA per day ([average MVPA min/day of weekdays × 5 + average MVPA min/day of weekend × 2] / 7).

PA was measured in 45 participants, and AF was measured in 43 participants. For eight participants (one male and seven females), AF values were missing due to absence on the day of the test, while for six participants (three males and three females), PA values were missing due to an insufficient number of valid measurement days (two weekdays and one weekend day). All participants had at least one measure of PA or AF.

### 2.3. Stimuli and procedure in MEG measurements

The cross-modal attention task (see Fig. 1) was programmed and controlled using PsychoPy software (Peirce et al., 2019). The participants were presented with continuous and simultaneous visual and auditory stimulation to produce frequency-tagged brain activity in each modality. The visual stimulus was varying visual noise, for which the luminance value of each pixel varied randomly between 0 and 255 at a frequency of 15 Hz, serving as the “tagging frequency” for the visual modality (adapted from Parkkonen et al., 2008). The auditory stimulus was a white noise stimulus that was amplitude-modulated at 40 Hz, serving as the “tagging frequency” for the auditory modality (adapted from Lamminmäki et al., 2014). An active task was used to guide the participants’ attention to one modality or another. The visual target was a square of 40 × 40 pixels for which the luminance value of each pixel varied randomly between 0 and 255, which appeared either in the left or right hemifield. The auditory target was a simple tone of 40 Hz delivered to either the left or right ear. The stimulation was divided into 36 separate blocks.

**Fig. 1.**
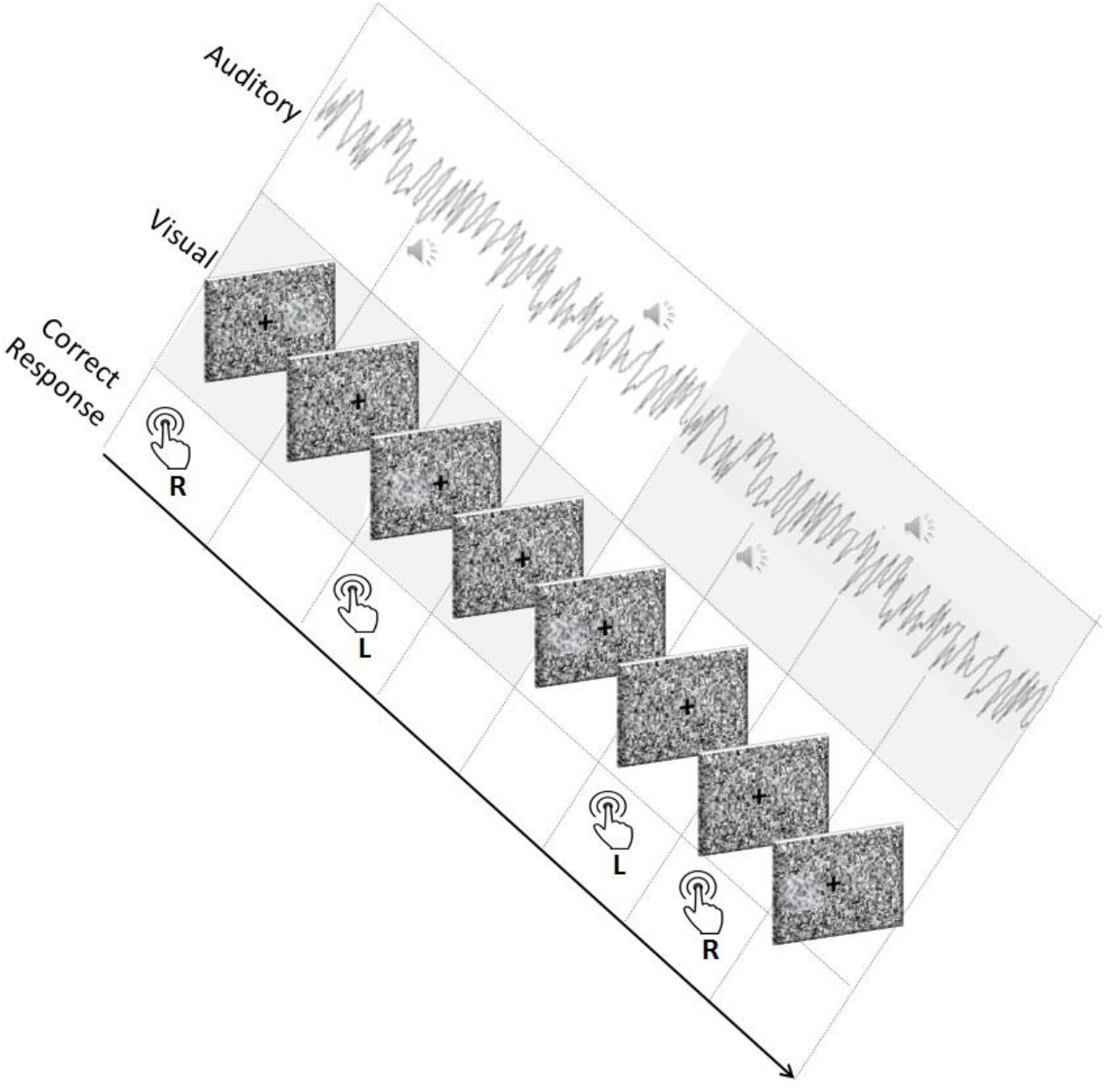
Schematic illustration of the progress of the cross-modal attention task. Dynamically varying noise at different frequencies for the visual and auditory modalities was embedded with left vs. right hemifield targets in each modality. At the beginning of each block, visual or auditory cues signaled whether attention should be directed toward visual or auditory modalities (attended modality shaded in gray color). Participants needed to indicate whether the target appeared on the left or right by pressing the corresponding button.

The duration of each block was approximately 36.5 seconds. Both auditory and visual target stimuli were present in each block, and a visual or auditory cue at the beginning of each block indicated which modality should be attended (nine blocks for visual and nine for auditory stimuli in a counterbalanced order). The cue indicating focusing on the visual modality was a white screen shown for a few seconds. The cue indicating focusing on the auditory modality was a tone with a frequency of 440 Hz. Each block contained eight randomly selected stimuli (four visual and four auditory) with interstimulus intervals of 4 s. Left and right stimuli had the same probability of occurrence.

After 18 blocks were administered, there was a break during which the participants could rest for some minutes. When the participants reported being ready to continue, the other 18 blocks were presented. Participants were instructed to report on which side (left or right) the target (visual or auditory stimuli, depending on the block) appeared by using MEG compatible response pads, and to ignore the stimulation modality not signaled at the beginning of each block. The total duration of the task was approximately 22 minutes.

To ensure that the participants had learned the instructions correctly, a practice session was performed prior to the real measurements. The practice lasted until the researchers considered that the adolescents had understood the task requirements. The participants understood the task only after each participant correctly identified at least two visual and two auditory targets on each side.

### 2.4. MEG recordings and analysis

Brain activity related to the cross-modal attention task was recorded using an Elekta Neuromag Triux system (MEGIN Oy, Helsinki, Finland). During the subjects’ preparation, five head-position indicator coils were attached to their heads in different locations (three on the frontal part of the head and one behind each ear). Coil’s locations were registered using a 3-D digitizer in relation to three anatomical landmarks (nasion and pre-auricular points). At the beginning of the recording, the participants’ position with respect to the helmet was measured and continuously tracked during the whole measurement (Uutela et al., 2001). Additional points located all around the head were also digitized. Eye movements were monitored using electro-oculograms (EOG), with two electrodes attached to the external canthi of both eyes for horizontal movements and two electrodes attached above and below the right eye for vertical movements. After the preparation period, the subjects were seated comfortably inside a magnetically shielded room and were instructed to avoid movement of the head and eyes. Two response pads (left and right) were positioned on a table attached to the chair and located over the subject’s legs. The task was projected onto a panel located one meter from the subject’s eyes.

MEG signals were band-pass filtered to 0.03–330 Hz and sampled at 1,000 Hz. The raw data were pre-processed using Maxfilter™ 2.2 software (MEGIN Oy, Helsinki, Finland). The signal space separation method (SSS) (Taulu et al., 2004) was used to remove the external interference emerging during the measurement. SSS was replaced by the spatiotemporal signal space separation method (tSSS) (Taulu & Simola, 2006) (also included in Maxfilter 2.2 software) for the analysis of data from participants wearing braces or other internal magnetic sources. Possible head movements occurring during the measurement with respect to the initial head position were also corrected using Maxfilter™ 2.2 software. The rest of the pre-processing was performed with Meggie (CIBR, Jyväskylä, Finland; Heinilä & Parviainen, 2022), a graphical user interface for MNE-Python (Gramfort et al., 2014). With the use of this software, epochs contaminated with eye movements (as measured with EOG) or cardiac artifacts (measured by an MEG channel) were removed from the analysis. Additionally, the data were visually inspected to exclude epochs contaminated by other kinds of artifacts.

The rest of the analysis was performed using Matlab (Mathworks Inc., 2020b). Analysis was performed with the brain activity recorded by planar gradiometers. Frequency spectra were computed with Welch’s method (Hanning window and FFT size 8,192 samples, 50% overlapping windows, frequency resolution 0.1221 Hz). Three ROIs consisting of 16 gradiometers (eight pairs), each located over the bilateral temporal cortex and the middle occipital cortex, were determined and used to compute the areal averages of the MEG signals. Identification of the visual and auditory frequency-tagged responses was performed based on peak frequency and amplitude. The visual tag-related signal was identified as the highest peak between 14 Hz and 16 Hz in the occipital areal average (peak always at 15.015 Hz). The auditory responses (left and right) were identified as the biggest peak between 39 Hz and 41 Hz in the bilateral temporal areal averages. The amplitude values for each ROI (at 15 Hz for bilateral occipital and at ca. 40 Hz for left and right temporal ROIs) were used for further analysis. The six resulting variables (visually attended and auditory attended in bilateral occipital, left temporal, and right temporal areas) and their individual variations were inspected to identify outliers. Typical values (Z-scores) that exceeded 2.5 standard deviations were considered outliers and were substituted with the closest non-atypical value.

In addition to contrasting the amplitude of the tag-related signals in the attended vs. unattended condition in each cortical ROI, we calculated the attention effects directly for both modalities. In the visual modality, the effect of attention was calculated as the normalized difference between the amplitudes at 15 Hz for the attended and unattended conditions in the occipital ROI. In the auditory modality, the effect of attention was calculated as the normalized difference between the amplitudes at 40 Hz for the attended and unattended conditions in both the left and right temporal ROIs. In both cases, normalization was done by subtracting the unattended value from the attended value and dividing the outcome by the value from the attended condition. The attention effects in each ROI were used for correlation analysis with the behavioral results from the cross-modal attention task.

### 2.5. Behavioral performance analysis

During the recordings, behavioral responses to the cross-modal attention task (the MEG task) were collected and further analyzed. Responses with a response time of less than 250 ms (less than 2% of the trials) were removed from further analysis for being considered too short to reflect attentional processing. From the remaining valid trials, the accuracy for each modality was calculated as the percentage of correct responses (in auditory attended and visual attended subtasks). Misses (ignored targets in the attended condition) in the visual and auditory subtasks were considered as indexing failed top–down attentional control and interpreted as inattention errors. False alarms (answers to unattended stimuli) in the visual and auditory subtasks were considered an indexing failure of inhibitory control.

Participants’ behavioral skills were also measured outside MEG recordings. Speed of attention was assessed with the reaction time (RTI) task, which is a subtest in the Cambridge Neuropsychological Test Automated Battery (CANTAB) (Cambridge Cognition Ltd., 2006). The RTI task measures reaction time (RT) and, specifically, the speed of response toward an unpredictable target. During the unpredictable condition, a yellow spot appeared in any of the five circles on the screen. Participants were instructed to retain their answers until they saw the yellow spot. Only then should they have touched, as fast as possible, the correct circle on the screen where the target had appeared. The subjects performed rehearsal trials until they understood the task and, afterward, 15 task trials. Scores were based on their RT (ms) and in-movement time (ms).

The participants’ inhibitory control skills were assessed using a modified Eriksen Flanker task (Eriksen & Eriksen, 1974). In each trial, the target (central arrow) was flanked by non-target stimuli (surrounding arrows). During the compatible condition, participants needed to report as fast as possible the target’s direction (left or right). During the incompatible condition, participants needed to report as fast as possible the opposite direction of the target (left button for the right direction of the target and right button for the left direction of the target). In both conditions, congruent trials (the target pointing in the same direction as the non-targets) and incongruent trials (the target pointing in the opposite direction to the non-targets) were included with the same probability of appearance. Accuracy (in percentages) and RT (in ms) were recorded separately for congruent and incongruent trials for each condition.

In the two additional behavioral tasks (Flanker and RTI), the variables regarding accuracy were removed from the final analysis due to reaching ceiling effects (more than 50% of the participants’ accuracy values were near the upper limit of the task range).

The resulting variables used for further analysis per task were as follows: (1) MEG task: Cross-modal attention task (misses for visual and auditory subtasks; false alarms for visual and auditory subtasks); (2) Flanker task (RT for compatible and incompatible conditions; error rates for incongruent trials); and (3) RTI task (RT for five-choice condition, movement time for five-choice condition). All the resulting behavioral variables were inspected to identify outliers. The values (Z scores) that exceeded 2.5 standard deviations were considered outliers (atypical) and were changed to the closest non-atypical values.

### 2.6. Statistical analysis

Statistical analysis was performed with SPSS (IBM SPSS Statistics for Windows, Version 26.0; IBM Corp., Armonk, NY, USA). To answer our research questions on the effect of attention and possible interaction with PA/AF in the engagement of sensory cortices, we conducted two repeated measures analyses of variance (rANOVA) on the neural engagement of visual and auditory cortices (strength of SSRs), one for each between-subjects factor (PA and AF groups). Within-subjects’ factors were ROI (occipital, left temporal, and right temporal) and power of SSRs based on task requirements (attended, unattended). Post-hoc comparisons tested the differences in estimated marginal means with Bonferroni correction for multiple comparisons.

Bivariate Spearman correlation coefficients with false discovery rate (FDR) corrections were used to test the associations between the attention effects (calculated as the normalized difference between attended and unattended conditions) at the visual and auditory sensory cortices and the behavioral performance of the cross-modal attention task (misses and false alarms in the visual and auditory subtasks).

One-way ANOVA was used to test differences between the high vs. low PA and AF groups in the behavioral (MEG) task and additional behavioral tasks.

## 3. Results

### 3.1. General spectral results

The existence of neural entrainment in the occipital and (left and right) temporal ROIs was confirmed for each participant. Figure 2 shows the power spectral density (PSD), evidencing a peak at around 40 Hz for the left and right temporal ROIs and at 15 Hz for the occipital ROIs. All participants showed 15-Hz SSRs (peak) in bilateral occipital ROIs (see Fig. 2C). However, only 35/51 participants showed 40-Hz SSRs in the left temporal ROI, and 36/51 participants showed 40-Hz SSRs in the right temporal ROI (see Fig. 2A, B). For comparison, the grand average PSD was plotted for individuals who showed responses, contrasted with those individuals who did not show responses.

**Fig. 2.**
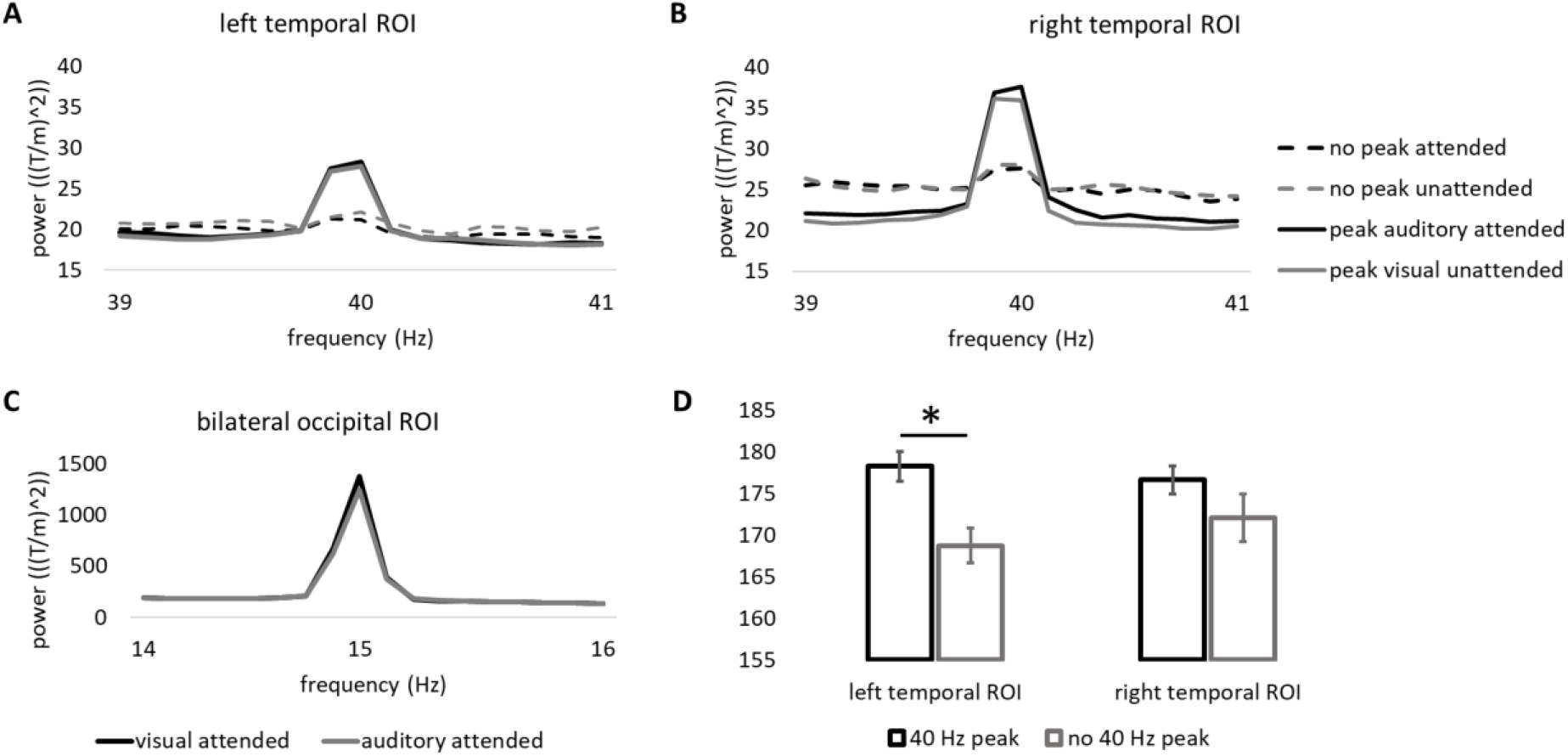
PSD showing neural engagement in visual and auditory primary sensory cortices for participants with and without tag-related responses. (A) PSD of the participants with (solid lines) and without (dashed lines) the 40-Hz peak for the attended (black) and unattended (gray) conditions in the left temporal area. (B) PSD of the participants with (solid lines) and without (dashed lines) the 40-Hz peak for the attended (black) and unattended (gray) conditions in the right temporal area. (C) PSD of the participants with the 15-Hz peak for the attended (black line) and unattended (gray line) conditions in the visual modality in the bilateral occipital areas. (D) Bar representation of the differences in age (months) for the participants with (black color) and without (gray color) 40-Hz SSRs in the left and right temporal ROIs.

To investigate whether the likelihood of SSRs at the individual level could indicate a state of maturation, we further tested the existence of age differences between participants with and without responses in both left and right temporal cortices with one-way ANOVA. Participants with 40-Hz SSRs in the left temporal ROI were older (178.28 ± 10.36 months) than participants without 40-Hz SSRs (168.75 ± 8.31 months) (F _(1, 50)_ = 10.44, *p* = 0.002). However, in the right temporal ROI, the age of participants with 40-Hz SSRs (176.64 ± 10.32) did not significantly differ from the age of participants without 40-Hz SSRs (172.07 ± 11.13, F _(1, 50)_ = 1.98, *p* = 0.165; see Fig. 2D).

### 3.2. Effects of attention on sensory cortices and their interaction with PA/AF in the brain

Both rANOVAs (PA and AF as between-subjects factor) showed the main effect of attention (PA as between-subjects factor: F _(2, 1.000)_ = 36.79, *p* < 0.001; AF as between-subjects factor (: F _(1, 1.000)_ = 29.95, *p* < 0.001). Both rANOVAs also showed a significant interaction of ROI x attention (PA as between-subjects factor: F _(2, 1.006)_ = 37.29, *p* < 0.001); AF as between-subjects factor: F _(2, 1.004)_ = 29.51, *p* < 0.001). Post-hoc comparisons showed that in the occipital ROI, the power of the 15-Hz SSRs was higher in the attended (1470.12 ± 149.03) condition than in the unattended (1338.48 ± 138.61) condition (*p* < 0.001). In the left temporal ROI, no differences were found between the attended (23.96 ± 1.51) and unattended (24.82 ± 1.77) conditions (*p* = 0.31). In the right temporal ROI, no differences were found between the attended (30.68 ± 2.32) and unattended (31.21 ± 2.68) conditions (*p* = 0.68).

Both rANOVAs showed a main effect of ROI (PA as between-subjects factor: F _(2, 1.000)_ = 92.47, *p* < 0.001; AF as between-subjects factor: F _(2, 1.000)_ = 105.07, *p* < 0.001) in SSR engagement, indicating stronger amplitudes for occipital than for left and right temporal areas. A between-subjects effect of group was found for PA (F _(1, 43)_ = 4.95, *p* = 0.03), but not for AF (*p* = 0.60). When PA was used as the between-subjects factor, a significant interaction between ROI × PA was found (F _(2, 1.000)_ = 5.24, *p* = 0.03). Post-hoc comparisons showed that the strength at the 15-Hz SSRs in the occipital ROI was higher (*p* = 0.03) for the participants in the high PA group (1729.73 ± 238.22) than for the participants in the modlow PA group (1078.87 ± 160.09) (see Fig. 3). No group differences were found for the left (*p* = 0.20) or right (*p* = 0.34) temporal ROIs. When AF was used as a between-subjects factor, no significant interaction of ROI x AF was found (*p* = 0.60).

**Fig. 3.**
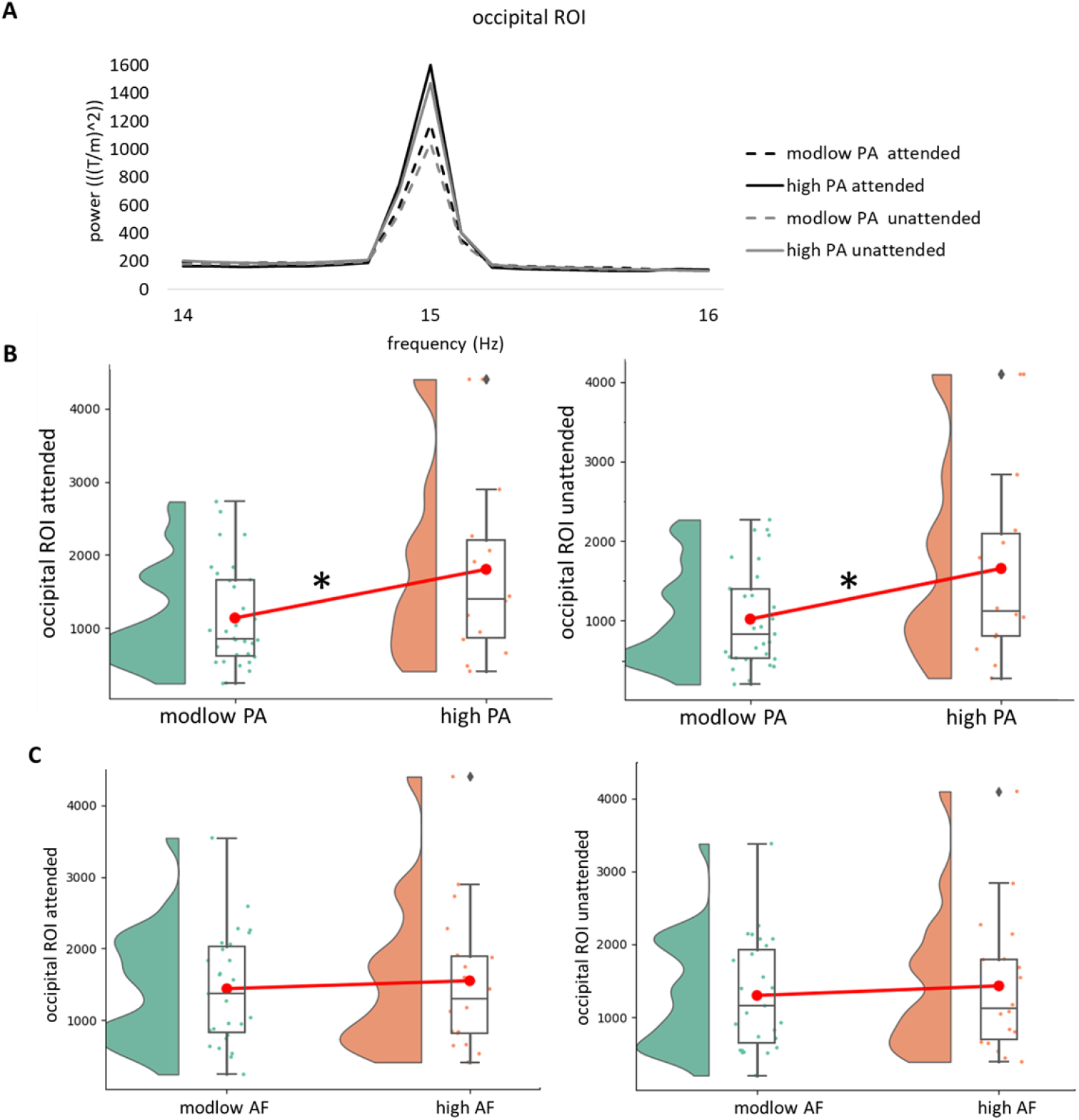
Modulation of SSRs in the bilateral occipital ROI for the PA groups. (A) Spectra representing the 15 Hz SSRs for the modlow (dashed lines) and high PA (solid lines) groups in the visual attended (black) and auditory attended (gray) conditions. (B) Raincloud plots showing the distributions and differences in the strength of the responses (attended and unattended) in the bilateral occipital ROI for the modlow (green) and high (orange) PA groups. (C) Raincloud plots showing the distributions and differences in the strength of responses (attended and unattended) in the bilateral occipital ROI for the modlow (green) and high (orange) AF groups.

### 3.3. Associations between brain measures and behavioral performance measures

A higher attention effect (i.e., increased amplitude for attended vs. unattended, calculated as the ratio between attended–unattended/attended) in the left temporal ROI was associated with more false alarms in the visual subtask (rho = 0.32, *p* = 0.02), although when FDR correction was applied, this correlation was not significant (*p* = 0.23). When the sample was divided by groups based on the presence of 40-Hz SSRs in the left temporal ROI, this correlation was significant for the group that did not show SSRs (rho = 0.54, *p* = 0.03) but not for the group that showed SSRs (*p* = 0.36) (see Fig. 4). No other significant correlations were observed for the cross-modal attention task with brain measures (attention effect occipital ROI vs. misses auditory p = 0.66, misses visual p = 0.47, false alarms auditory p = 0.66, false alarms visual p = 0.40; attention effect left temporal ROI vs. misses auditory p = 0.16, misses visual p = 0.99, false alarms auditory p = 0.29; attention effect right temporal ROI vs. misses auditory p = 0.74, misses visual p = 0.54, false alarms auditory p = 0.66, false alarms visual p = 0.36).

**Fig. 4.**
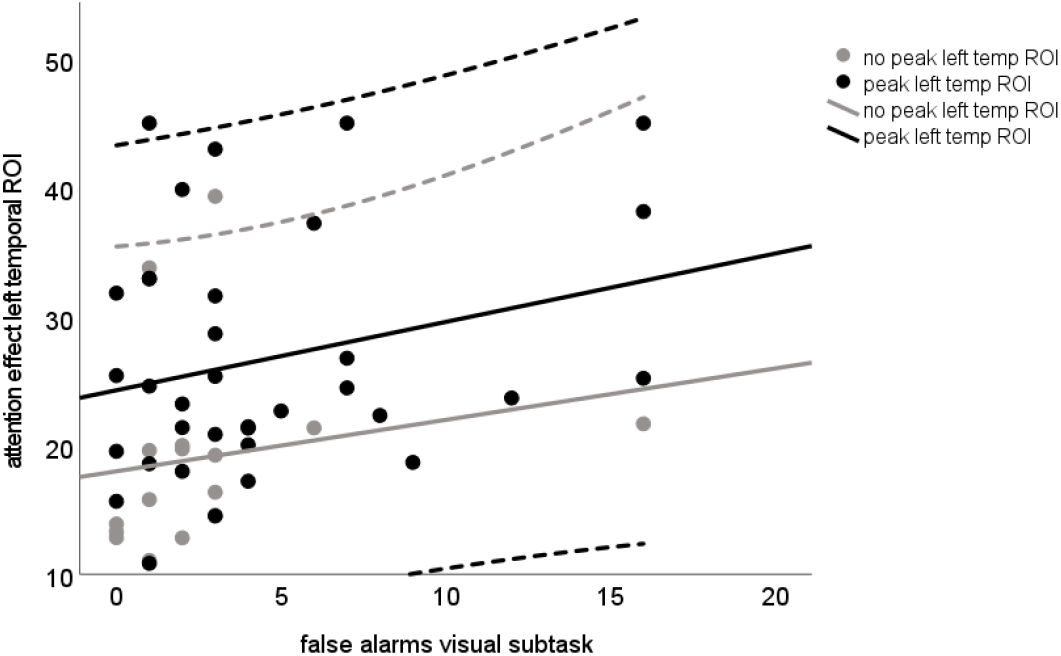
Association between the attention effect in the left temporal ROI and the number of false alarms in the visual subtask. The black color represents the participants with 40 Hz SSRs in the left temporal ROI. Gray represents the adolescents without 40 Hz SSRs in the left temporal ROI. The solid central lines represent the linear trend line, and the two external dashed lines represent the 95% confidence interval.

No significant correlations were found for RTI and Flanker tasks with brain measures (attention effect occipital ROI vs. RTI RT p = 0.14, RTI movement time p = 0.17, Flanker congruent RT p = 0.72, Flanker incongruent RT p = 0.90, Flanker incongruent false alarms p = 0.48; attention effect left temporal ROI vs. RTI RT p = 0.10, RTI movement time p = 0.49, Flanker congruent RT p = 0.59, Flanker incongruent RT p = 0.80, Flanker incongruent false alarms p = 0.49; attention effect right temporal ROI vs. RTI RT p = 0.06, RTI movement time p = 0.93, Flanker congruent RT p = 0.26, Flanker incongruent RT p = 0.38, Flanker incongruent false alarms p = 0.49).

### 3.4. Behavioral performance in cross-modal attention (MEG) task, RTI and Flanker tasks, and their association with physical activity and aerobic fitness

The behavioral performance in the cross-modal attention task of participants in the high PA group did not differ from that of participants in the modlow PA group (misses visual subtask p = 0.91, misses auditory subtask p = 0.13, false alarms visual subtask p = 0.59, false alarms auditory subtask p = 0.28) (see Fig. 5A). Similar results were obtained for the additional behavioral tasks, RTI (mean five-choice RT p = 0.93, mean five-choice movement time p = 0.52), and Flanker tasks (RT compatible condition p = 0.94, RT incompatible condition p = 0.82, errors incongruent trials p = 0.41) across groups of PA (see Fig. 5B).

**Fig. 5.**
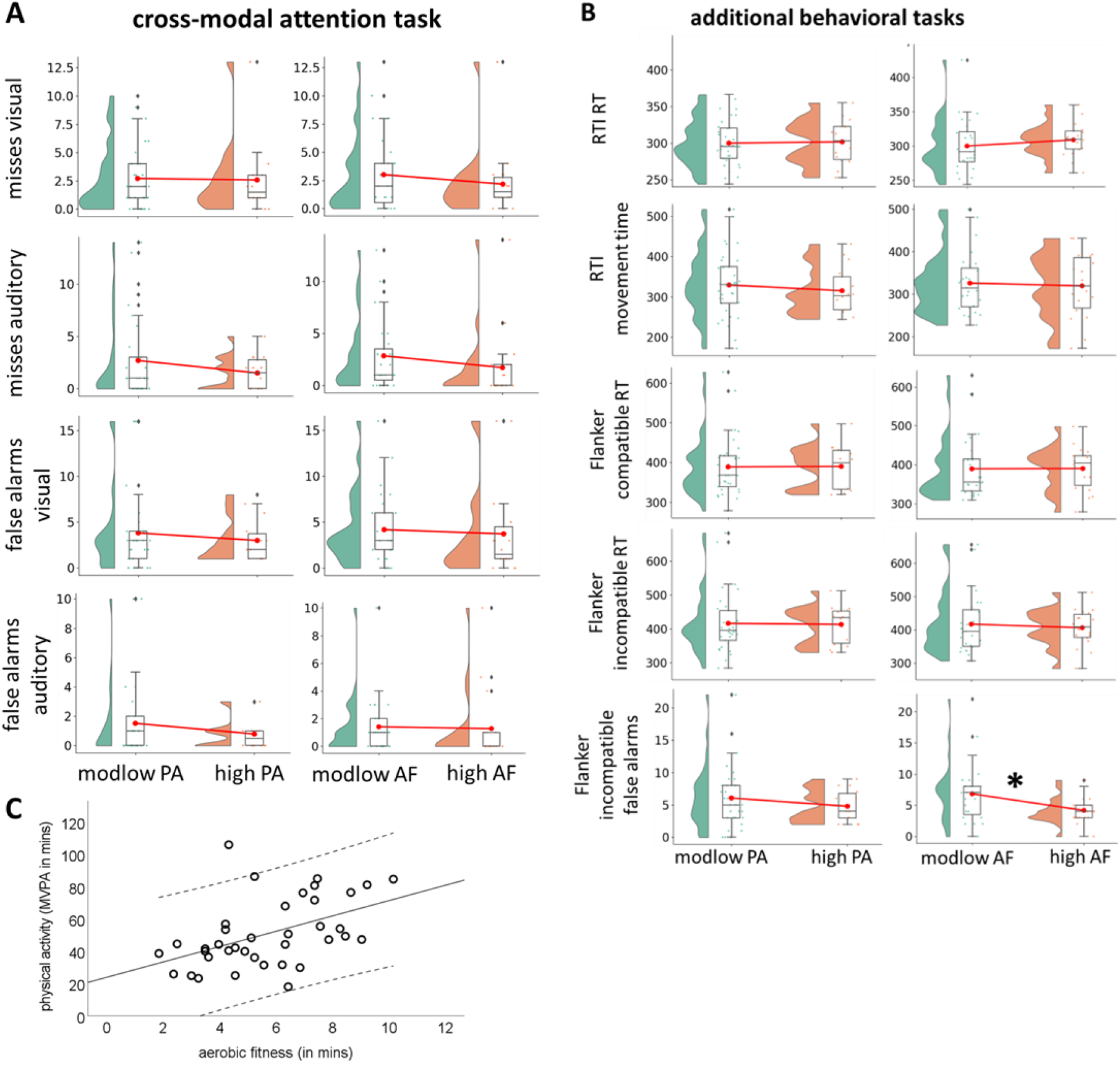
Raincloud plots showing the distribution of variables for modlow (green) and high (orange) groups of physical activity and aerobic fitness (solid red line connects the groups’ mean value) in (A) the cross-modal attention task and (B) the additional behavioral tasks (RTI and Flanker). (C) Scatterplot showing the association between physical activity and aerobic fitness.

The behavioral performance in the cross-modal attention task of participants in the high AF group did not differ from that of participants in the modlow AF group (misses visual subtask p = 0.41, misses auditory subtask p = 0.24, false alarms visual subtask p = 0.79, false alarms auditory subtask p = 0.78) (see Fig. 5A). Similar results were obtained in the RTI test (mean five-choice RT p = 0.43, mean five-choice movement time p = 0.71). Adolescents in the high AF group made fewer errors (4.17 ± 2.12) in the incongruent trials of the Flanker task than those in the modlow AF group (6.72 ± 4.95) (t [1, 34.611] = 2.30, p = 0.03); see Fig. 2A. RTs in the compatible and incompatible conditions of the Flanker task did not differ across the AF groups (RT-compatible p = 0.89, RT-compatible p = 0.57) (see Fig. 5B).

There was a significant correlation between PA and AF (rho = 0.524, p < 0.001; see Fig. 5C).

## 3. Discussion

We explored the neuromagnetic indicators of cross-modal attention, namely frequency-tagged SSRs in visual (occipital) and auditory (temporal) cortices during audiovisual task performance, and their link with behavioral performance. Second, we clarified that the level of PA, but not AF, is associated with the brain basis of cross-modal attention, especially in visual areas, in typically developing adolescents.

Contrary to our expectations, we found divergent engagement of visual versus auditory sensory cortices during the cross-modal attention task. Bilateral occipital cortices showed clear SSRs for visual stimulation at 15 Hz in all participants. This general engagement of visual areas suggests that the primary visual cortical areas follow dynamic changes in sensory stimulation with the presented temporal frequency. However, the expected SSRs for auditory stimulation at 40 Hz in the bilateral temporal cortices were only observed in 35 out of 51 participants for the left temporal cortex, and 37 participants in the right temporal cortex.

The less systematic response of auditory areas could be explained by the developmental stage of the adolescent brain. During brain development, synaptic pruning (an experience-dependent loss of synaptic connections) occurs at separate times in different cortical areas. The visual cortex has been suggested to reach maturity during childhood by about 7 years of age (Huttenlocher, 1990), while the auditory cortices mature later during adolescence (Gogtay et al., 2004; Ponton et al., 2000) by about 12 years of age (Huttenlocher & Dabholkar, 1997). Indeed, earlier neuromagnetic studies clearly show immature auditory responses in children at the age of 7 years (Parviainen et al., 2019) that persist at least until the age of 13 (Sussman et al., 2008; van Bijnen et al., 2022). For the cortical circuits to be able to generate SSRs for external acoustic information, especially at the high frequency range of 40 Hz, the synaptic connections need to have reached the needed temporal precision. It is exactly the temporal processing properties that have been suggested to demonstrate a protracted developmental trajectory in cortical circuits (Ponton et al., 2000).

Another possible explanation is that the different tagging frequencies between the cortices (15 Hz for occipital ROI and 40 Hz for left and right temporal ROIs) could underlie these effects. Indeed, it may be that cortical engagement is more pronounced at lower frequencies in any sensory processing (Albouy et al., 2022; Otero et al., 2022). Particularly in auditory areas, the importance of lower frequencies for language processing in the brain has been highlighted (Haegens & Golumbic, 2018), specifically for processing words (Kolozsvári et al., 2021) and sentences (Kolozsvári et al., 2021; Molinaro & Lizarazu, 2018). However, the frequencies used in the current study were chosen based on earlier literature that showed an increase in the entrainment of oscillatory activity in the human auditory cortex triggered by about 40 Hz stimulation frequencies (Lamminmäki et al., 2014; Ross et al., 2005), as in the case of auditory steady-state responses (ASSRs). Even though ASSRs are known to be generated in the brainstem by using animal models, the generators of 40-Hz ASSRs have also been localized in the primary auditory cortex (Li et al., 2018).

In line with the maturational hypothesis, the adolescents with 40 Hz SSRs were older than those who did not show SSRs, but only for the left temporal area. Our findings are in line with those studies suggesting that the maturation of the right hemisphere auditory cortex might precede the maturation of the left hemisphere (Edgar et al., 2016; Herdman, 2011; Ono et al., 2020; Parviainen et al., 2011, 2019). More studies with longitudinal follow-up measurements would be needed to reveal possible maturational differences in engagement across hemispheres in adolescents. In general, our findings provide evidence of a developmental effect on the already known asymmetric organization of the auditory primary cortices at the level of sensory processing (Hernández et al., 2022; Parviainen et al., 2005; Poeppel, 2003).

We expected that SSRs in both visual and auditory sensory cortices would show increased amplitude in the attended condition compared to the unattended condition. This hypothesis was confirmed for the visual cortex, where the effect of attention was observed. In this regard, the results are in line with those studies showing attention-induced changes in sensory cortices in unimodal (Fiebelkorn et al., 2018) and multimodal tasks (Plöchl et al., 2022). In sum, our findings partially support the idea of modulation of neural activity in the sensory cortices exerted by top–down attentional control, as that modulation was only found for the visual cortex in our study.

No effect of attention was observed in the auditory cortices of the SSRs. This can be linked with the developmental effects found in auditory cortices (i.e., the 40 Hz stimulation did not lead to observable responses in ca. 30% of the subjects). If the cortical circuits fail to show SSRs with the (bottom–up) representation of high-frequency acoustic modulation in the sound, it may also be that the connections with the areas providing top–down modulation to the sensory circuits are not yet fully in function. Perhaps the expected attentional modulation can be seen later in the lifespan. Future studies comparing adolescents and adults are needed to disentangle the developmental influence from other neuronal mechanisms that might be underlying the cross-modal induced changes in sensory cortices.

As expected, based on earlier literature, only PA, but not AF, showed an association with cortical processes during the cross-modal attention task. Our results showed higher neural engagement of bilateral occipital cortices during attended and unattended conditions for participants with high PA than for those with moderate to low PA, while no effects for AF were found. One possible contributing factor is that the above-mentioned maturational effects are involved in the increased neural engagement of bilateral occipital cortices for adolescents with higher levels of PA. During adolescence, synaptic connections that do not participate in behaviorally meaningful functional networks and neural coding are pruned. This is especially true for occipital areas (Rauschecker & Marler, 1987). It may be that individuals with higher levels of PA are more exposed to visual material in general, and thus, visual areas are more strongly functionally coupled with sensory input during the first two decades of life. This, however, seems to be a purely sensory/automatic process, as there was no difference between the attended and unattended conditions. In general, our results support the idea that PA might be related to functional changes in the brains of adolescents more strongly than AF, as suggested by previous studies from our research group (Hernández et al., 2021; Ruotsalainen et al., 2020; 2021). More developmental studies approaching this issue comparing adolescents with children and adults are needed to confirm the contribution of maturational changes to this effect.

Higher-fit participants showed fewer errors in the additional Flanker task. This finding supports previous studies reporting better inhibitory skills associated with higher AF in adolescents (Huang et al., 2015; Shigeta et al., 2021; Stroth et al., 2009; Westfall et al., 2018). However, AF (as well as errors in the Flanker task) were not linked to the neuromagnetic signatures of cross-modal attention in the visual and auditory cortices. This suggests that AF might be involved with inhibitory control processes in adolescents, but this effect does not seem to be related to their brain processing of cross-modal attention in sensory cortices to the extent that we probed it with the SSRs.

PA was not associated with any behavioral effect with measures from Flanker, RTI, or cross-modal tasks. Adolescents with moderate to low PA performed the cross-modal attention task equally well as did the participants with high PA. This suggests that group differences in bilateral occipital activation patterns are not linked with performance differences, but rather reflect enhanced reactivity of the visual cortex during cross-modal task. Previously, structural changes in occipital regions have been associated with high PA levels, especially in older populations (Dawe et al., 2021; Erickson et al., 2010) and midlife adults (Tarumi et al., 2021). These changes were related to either increased cortical thickness due to gray matter volume (Erickson et al., 2010) or white matter properties (Dawe et al., 2021). Functional studies have also found links between higher PA levels and occipital brain areas related to pre-attentional processing (Pesonen et al., 2019) in adults. In adolescents and children, this kind of finding is rarer. However, in line with our results, Ludyga et al. (2018) reported a PA-related amplitude increase of visual P300 in parieto-occipital regions for 12- to 15-year-old adolescents. This amplitude increase was associated with better inhibitory control skills. Altogether, these findings suggest that PA might start influencing occipital areas around adolescence and that these changes remain during adulthood.

Thus, we provide evidence suggesting that visual and auditory cortices, as they have been studied here (by using SSRs in visual and auditory cortices), do not necessarily operate in a mutually exclusive way. This means that they may be part of a common neural network involved in cross-modal attention that, due to maturational reasons, seems to be led by bilateral occipital areas in adolescence.

As a controversial result, the 40-Hz SSRs in the left temporal ROI were linked with the cognitive performance level in the cross-modal attention task. Interestingly, this association was specifically driven by the participants without a recognizable peak at the 40 Hz frequency band. The increased baseline amplitude at 40 Hz (but not the SSRs) correlates positively with the failure to inhibit unattended stimuli (i.e., increment of false alarm responses; in other words, responses to stimuli in the unattended [visual] stream). This suggests that the level of engagement of the left auditory cortex during audiovisual cross-modal attention tasks might reflect the efficiency of inhibitory mechanisms in the brain. It might be that the lack of maturation in the left temporal cortex SSRs at 40 Hz during adolescence is behaviorally expressed as deficient inhibitory control during cross-modal processing, but this was not tested in our study. These results are consistent with previous studies showing a link between increased activation in temporal areas and high load inhibitory processing in typically developed adolescents (McAlonan et al., 2009; Woolley et al., 2008). Perhaps, at the early stages of development, a compensatory strategy of the still immature left temporal cortex to deal with cross-modal attention is to increase 40 Hz responses to relevant (auditory) information at the expense of not successfully inhibiting responses to irrelevant stimuli.

A possible neurophysiological explanation for this effect could be related to the role of gamma band oscillations in the brain. Despite the 40 Hz SSRs in our study, gamma band oscillations and 40 Hz ASSRs are distinct phenomena; they might share some properties because they all oscillate at a similar frequency. Cognitive gamma oscillations (40–120 Hz) have been previously identified as supporting attention/inhibitory processing (Fries et al., 2001), and inhibitory GABAergic interneurons seem to be part of their key circuitry (Wang & Buzsáki, 1996). Indeed, auditory 40 Hz ASSRs/gamma oscillations have been associated with inhibitory control abnormalities in disorders such as autism (Seymour et al., 2020) and schizophrenia (Metzner et al., 2022). This also suggests that the auditory 40 Hz SSRs in the left temporal cortex in the present study, and its age effects in adolescents, could reflect the developmental stage of inhibitory processes in these cortical networks.

There are some limitations to this study. First, due to the lack of 40 Hz SSRs observed for a group of adolescents in the left and right temporal areas, the effect of attention was not equally reliably approachable for the auditory than for the visual domain. Another limitation of this study is related to the relatively small sample size. When treated as one group, the associations of PA/AF levels were approachable with the present sample, but the individual variance in the strength of the responses weakened further comparisons. With a larger sample, it would have been possible to split the sample based on the presence of SSRs in the auditory cortices.

In future studies, it would be important to investigate these processes in a small age range while ensuring a large enough sample size, given the strong variation of the auditory cortex signaling shown within our sample. Finally, we focused our methodological approach on attentional effects at the sensory level of processing in the visual and auditory cortices, which is a new approach, especially in developmental brain stages. Naturally, other brain areas are known to contribute to the top–down control of cross-modal attention, and our results must be seen as complementing the existing literature that so far has contributed more to uncovering the role of the “higher level” association areas.

## 5. Conclusion

The brain network underlying audiovisual cross-modal attention processing in the adolescent brain shows unequal involvement of visual and auditory sensory cortices. All of the adolescents showed the expected tag-driven signals in the primary visual cortex, while primary auditory cortices showed partial involvement in the cross-modal task, with more immature involvement of the left auditory cortex. These findings suggest that there are developmental differences between auditory and visual sensory cortices in temporal precision during adolescence. The visual sensory cortex showed enhanced tag-related activity with attention. The steady-state responses in the occipital cortex were enhanced in adolescents with higher physical activity but not with higher aerobic fitness, irrespective of task demands.

## Acknowledgements

The authors would like to thank Jaakko Leppäkangas for his valuable help in coding the cross-modal attention task. MEG recordings were conducted with the support of the Academy of Finland, grant numbers 273971, 274086, and 311877. This study was also supported by a personal grant to DH from the Jenni and Antti Wihuri Foundation. We would like to thank Josh Seligman for the English Language proofreadings of this manuscript.

